# Frequent Aneuploidy in Primary Human T Cells Following CRISPR-Cas9 cleavage

**DOI:** 10.1101/2021.08.20.457092

**Authors:** A.D. Nahmad, E. Reuveni, E. Goldschmidt, T. Tenne, M. Liberman, M. Horovitz-Fried, R. Khosravi, H. Kobo, E. Reinstein, A. Madi, U. Ben-David, A. Barzel

## Abstract

Multiple ongoing clinical trials use site-specific nucleases to disrupt T cell receptor (TCR) genes in order to allow for allogeneic T cell therapy^1–5^. In particular, the first U.S. clinical trial using CRISPR-Cas9 entailed the targeted disruption of the TCR chains and programmed cell death protein 1 (PDCD1) in T cells of refractory cancer patients^6^. Here, we used the same guide RNA sequences and applied single-cell RNA sequencing (scRNAseq) to more than 7000 primary human T cells, transfected with CRISPR-Cas9. Four days post-transfection, we found a loss of chromosome 14, harboring the TCRα locus, in up to 9% of the cells, and a chromosome 14 gain in up to 1.4% of the cells. We further identified truncations of chromosome 7, harboring the TCRβ locus, in 9.9% of the cells. Loss of heterozygosity (LOH) was further validated using fluorescence *in situ* hybridization (FISH) and the temporal dynamics of cleavage and incomplete repair were monitored using digital droplet PCR (ddPCR). Aneuploidy was found among all T cell subsets and was associated with transcriptional signatures of reduced proliferation and metabolism as well as with induced p53 activation and cell death. We conclude that aneuploidy and chromosomal truncations are frequent outcomes of CRISPR-Cas9 cleavage in clinical protocols. Monitoring and minimizing these aberrant products is crucial for future applications of genome editing in T cell engineering and beyond.

## MAIN

Engineered T cells are revolutionizing cancer immunotherapy. Notably, four treatments entailing chimeric antigen receptor (CAR) T cells have received regulatory approval in different B cell malignancies^7^. At the same time, hundreds of ongoing clinical trials, for multiple additional indications, are applying different variations of CARs or engineered TCRs in an attempt to improve efficacy, safety and scalability^1^. Importantly, the approved CAR T therapies all entail the cumbersome, expensive and time-consuming manipulation of autologous T cells. Many current efforts are therefore directed at facilitating the engineering and banking of allogeneic CAR/TCR-T cell products. Importantly, the endogenous TCR of engineered allogeneic T cells has to be disrupted in order to prevent graft vs. host disease (GVHD) and this is most often achieved using site-specific nucleases such as meganucleases^2^, TALENs^3,4^, megaTALs and CRISPR-Cas9^5^. Cleaving the TCR loci can further serve for the directed integration of CAR and engineered TCR genes, allowing uniform expression, enhancing T cell potency and delaying exhaustion^8– 10^. Finally, additional genes, beyond the TCR chains, may be disrupted in order to improve T cell function^11^, confer drug resistance^12^, or avoid checkpoint inhibition^13^.

Site-specific nucleases can be highly potent^14^, but they are associated with a variety of undesired outcomes. Cas9 can be immunogenic *in vivo*, eliciting both humoral and cellular responses^15^. We recently demonstrated that Cas9 activates the p53 pathway and selects for p53-inactivating mutations^16^. Off-target cleavage can in turn be reduced, but not eliminated^17^, and small on-target insertions and deletions (InDels) are common, even when providing a donor template for gene correction or insertion. Importantly, CRISPR-Cas9 cleavage can also lead to gross chromosomal aberrations. In particular, CRISPR-Cas9 cleavage leads to large deletions in early mouse^18^ and human embryos^19^ as well as in embryonic stem cells and induced pluripotent stem cells^20^. Moreover, chromosomal truncations were reported following CRISPR-Cas9 cleavage in mouse and human cell lines^21,22^, and entire chromosome loss has resulted from CRISPR-Cas9 cleavage in human embryos^23^. Recently, chromothripsis, defined as multiplicity of gross genomic rearrangements in a one-off cellular crisis, was shown to occur in mobilized human CD34^+^ cells following CRISPR-Cas9 cleavage of the β-globin gene in a clinically relevant setting^24,25^, with possible ramifications for the development of gene therapies for hemoglobinopathies. Finally, in the first U.S. clinical trial involving CRISPR-Cas9, Stadtmauer & Fraietta et al. aimed at disrupting the TCR and PDCD1 loci for allogeneic T cell therapy and reported the detection of chromosomal translocations whose frequency decreased over time after infusion into patients^6^.

Here, we use the same guide RNA sequences, as did Stadtmauer & Fraietta et al.^6^, to target the TCR and PDCD1 loci in primary human T cells with CRISPR-Cas9. Using a novel unbiased approach, based on scRNAseq, corroborated by FISH and ddPCR analyses, we detect frequent aneuploidy and truncations of the chromosomes harboring the targeted loci.

We hypothesized that double-strand DNA breaks, induced by site-specific nucleases, may sometime fail to be repaired and result instead in chromosomal truncations and aneuploidy. We further conjectured that such adverse outcomes could be detected and monitored, in a high throughput manner, by following concerted changes in gene expression along chromosomes. In order to analyze chromosomal truncations and aneuploidy in a clinically relevant setting, we first performed CRISPR-Cas9 ribonucleoprotein (RNP) electroporation into primary human T cells using a single guide RNA (gRNA) targeting the TCRα locus (Fig. 1A). Importantly, we used the same gRNA sequence as used clinically in T cells by Stadtmauer & Fraietta et al.^6^. As a control, we used an irrelevant gRNA^14^, with no matching target in the human genome. The TCRα locus was successfully targeted in more than 50% of the cells (Fig. 1B-C, Extended Data Fig. 1). 4 days post-transfection, samples were subjected to scRNAseq (Methods). The transcriptional downregulation of TCRα was confirmed (Fig. 1D), and the transcriptional landscapes of the cells were used to infer copy number alterations using a well-established method (inferCNV)^26,27^. Strikingly, 5.3% of the cells in the TCRα targeted sample had expression patterns indicating a chromosome 14 loss (Fig. 1E-F). In addition, among these cells, chromosome 14 was an extreme outlier in the mean number of genes with undetected expression (p value< 0.0001, Extended Data Fig. 2). Notably, a chromosome 14 q-arm truncation at the TCRα locus is functionally equivalent to a whole-chromosome aneuploidy, because almost all protein-coding genes on chromosome 14 are coded on the q-arm, distal to the TCRα locus (Fig. 1A). Interestingly, a smaller fraction of the cell population, entailing 0.5-0.9% of the cells, had an apparent gain of chromosome 14 (Fig 1E-F). This functional chromosome 14 gain is assumed to result from mis-segregation of an acentromeric q-arm^24^. In particular, 100 chromosome 14 genes were underexpressed and 107 chromosome 14 genes were overexpressed in the cells categorized as harboring a chromosome 14 loss or gain, respectively (Extended Data Fig. 2, Supplementary Table 1). To set a clear statistical threshold for gains and losses, only cells with inferCNV scores >2 standard deviations from the mean score of the population were defined as having a chromosomal aberration (Methods).

**Figure 1:**
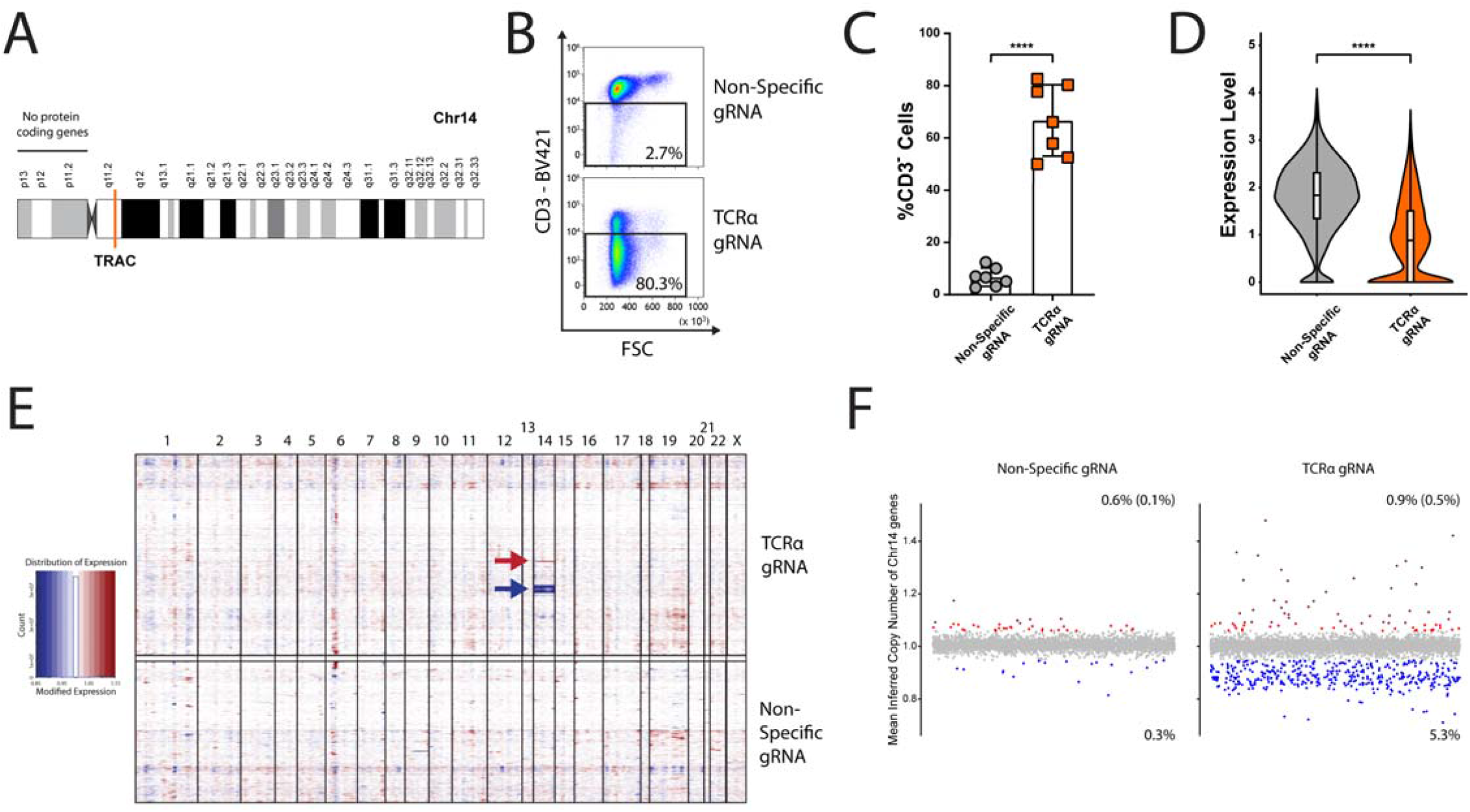
Targeting the TCRα locus using CRISPR-Cas9 leads to chromosome 14 aneuploidy. **A**. Schematic depiction of human chromosome 14 (Chr14). The target locus is indicated in orange. **B**. Flow cytometry example of TCR ablation in primary human T cells following CRISPR-Cas9 RNP electroporation. Cells were electroporated with Cas9 and either a non-specific gRNA or a TCRα-targeting gRNA. **C**. Quantification of B. Each dot represents an independent experiment. Mean value, standard deviation and individual experiments are indicated. n=7, ****, p<0.0001, two-sided unpaired Wilcoxon test. **D**. A reduction in TCRα expression is evident in the scRNAseq. The violin plots correspond to TCRα expression level in cells treated with either the non-specific gRNA or the TCRα-targeting gRNA. Upper and lower boundaries, median and quartiles are indicated. **** = p<0.0001, two-sided unpaired Wilcoxson test. **E**. A heat map depicting gene copy numbers inferred from scRNAseq analysis following treatment with either the non-specific or TCRα-targeting gRNAs. The Chromosomes are ordered in columns and the color coding indicates an increase (red) or decrease (blue) in copy number of genes along the chromosomes (x-axis) and each line represents an individual cell. Arrows indicate gain (red) or loss (blue) of chromosome 14. F. Each dot represents the mean inferred copy number of genes coded on chromosome 14 in each cell treated with a non-specific gRNA (left) or a TCRα-targeting gRNA (right). Cells are marked with red and blue dots, corresponding to a chromosome 14 gain or loss respectively, if their mean inferred gene copy number is >2 standard deviations (blue and red), or 3 standard deviations (dark red), from the population’s mean. n=6055 and 7700 for Non-specific (left) or TCRα-targeting (right) gRNA treated cells, respectively.

In order to corroborate the scRNAseq results, we next performed FISH assays employing red and green probes proximal and distal to the TCRα locus on chromosome 14, respectively. Expectedly, most examined nuclei had two foci of co-localized red and green signals (often co-detected as a yellow signal). Importantly, in two independent FISH iterations, signal patterns indicative of aneuploidy were observed significantly more frequently in TCRα-targeted cells than in control cells (Fig. 2A, Extended Data Fig. 3). These cells had either a single focus of co-localized signals or one focus of co-localized signals and another red focus, interpreted as corresponding to a truncation distal to the targeted gene. In particular, the 4% excess rate of these FISH patterns in TCRα-targeted cells, compared to control cells, is in agreement with the scRNAseq analysis.

**Figure 2:**
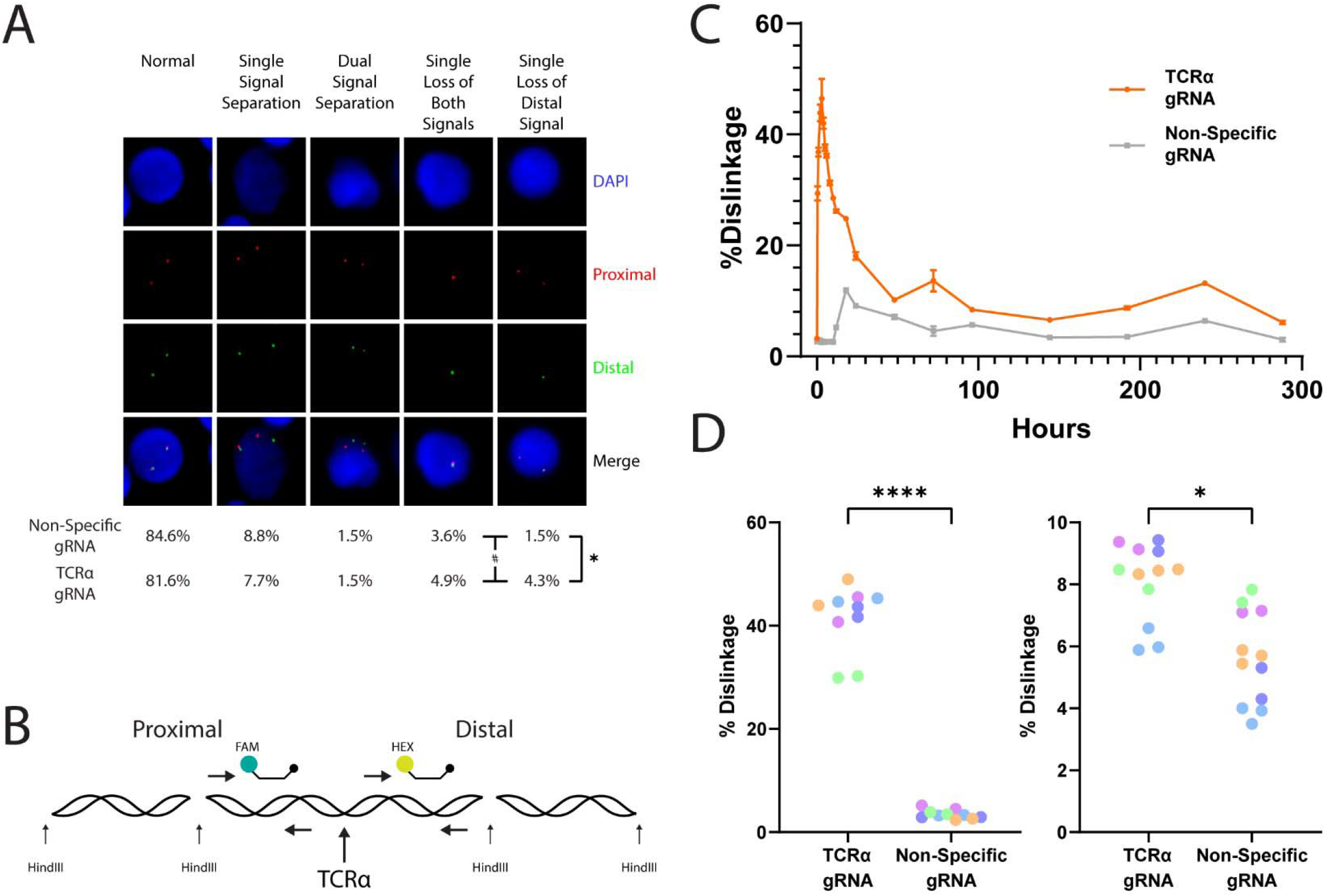
FISH and ddPCR analyses of Chromosome 14 aberrations. **A**. FISH analysis of cells treated with either a non-specific or a TCRα-targeting gRNA. In the pictures are examples of all signal patterns found among analyzed cells. The frequency of each signal pattern, among cells electroporated with the TCRα-targeting gRNA or a non-specific gRNA, is presented below the pictures. n = 273 and n = 506 for cells electroporated with the non-specific and TCRα-targeting gRNA, respectively. * p=0.0129 Fisher’s exact test for loss of distal signal comparison. #, p=0.0486 Fisher’s exact test for loss of either both signals or the distal signal. **B**. Schematic depiction of the ddPCR experiment. Genomic DNA is sheared using the HindIII restriction enzyme. The two sets of primers and probe are on the same restriction fragment are separated only upon CRISPR-Cas9 cleavage. **C**. Dislinkage, calculated based on the rate of single and dual color droplets (see methods) is analyzed at different time points following CRISPR-Cas9 electroporation with either a TCRα-targeting gRNA (orange) or a non-specific (gray) gRNA from a single experiment. Mean and standard deviation for technical replicates are indicated. **D**. Dislinkage at either 3 hours (left) or 4 days (right) following CRISPR-Cas9 electroporation. Each color represents a different human donor. Each dot represents a technical replication. n=5, ****, p<0.0001 and *, p=0.0277, two-sided unpaired t-test.

The rate of chromosome 14 breakage and its incomplete repair can further be monitored using digital droplet PCR (ddPCR)^28^ (Fig. 2B). DNA was collected at different time points after CRISPR-Cas9 RNP electroporation, in order to assess the dynamics of dislinkage between the two sides of the break. As early as 3 hrs after electroporation, the rate of dislinkage has peaked at more than 45% (Fig. 2C-D). Dislinkage rates were greatly reduced by 24 hrs, but remained significantly higher in the TCRα-targeted sample, compared to the control sample, for at least 4 days, indicating incomplete repair. While this ddPCR analysis alone cannot discriminate between transient breaks, translocations and deletions of various sizes, its results are consistent with the results from the scRNAseq and FISH analyses. Moreover, the ddPCR assay provides a scalable and cost-effective means to track the dynamics of breakage, repair and aberration.

We next used t-distributed stochastic neighbor embedding (t-SNE) analysis in order to cluster T cells according to their transcriptional signatures (Fig. 3A, Extended Data Fig. 4). We found a similar distribution, among the clusters, of T cells transfected with CRISPR-Cas9 RNPs entailing either the TCRα-targeting gRNA or an irrelevant gRNA with no human target (Fig. 3B). Moreover, among cells receiving the TCRα-targeting gRNA, neither cells with chromosome 14 loss nor cells with chromosome 14 gain were found to be considerably enriched in any T cell subset (Fig. 3C). This implies unbiased incidence of aneuploidy, as well as similar selection forces acting on aneuploid cells of various T cell states. Importantly, however, a gene set enrichment analysis (GSEA) detected significant differences in global gene expression patterns between the groups, beyond the expected reduced expression of the genes encoded on chromosome 14 (Fig. Extended Data Fig. 5). Specifically, the transcriptional signatures enriched in T cells with a chromosome 14 loss or gain reflected reduced expression of genes associated with the cell cycle and with various metabolic pathways, and increased expression of genes associated with p53 pathway activation and with apoptosis, implying reduced fitness of the aneuploid cells (Fig. 3D-F and Supplementary Table 2).

**Figure 3:**
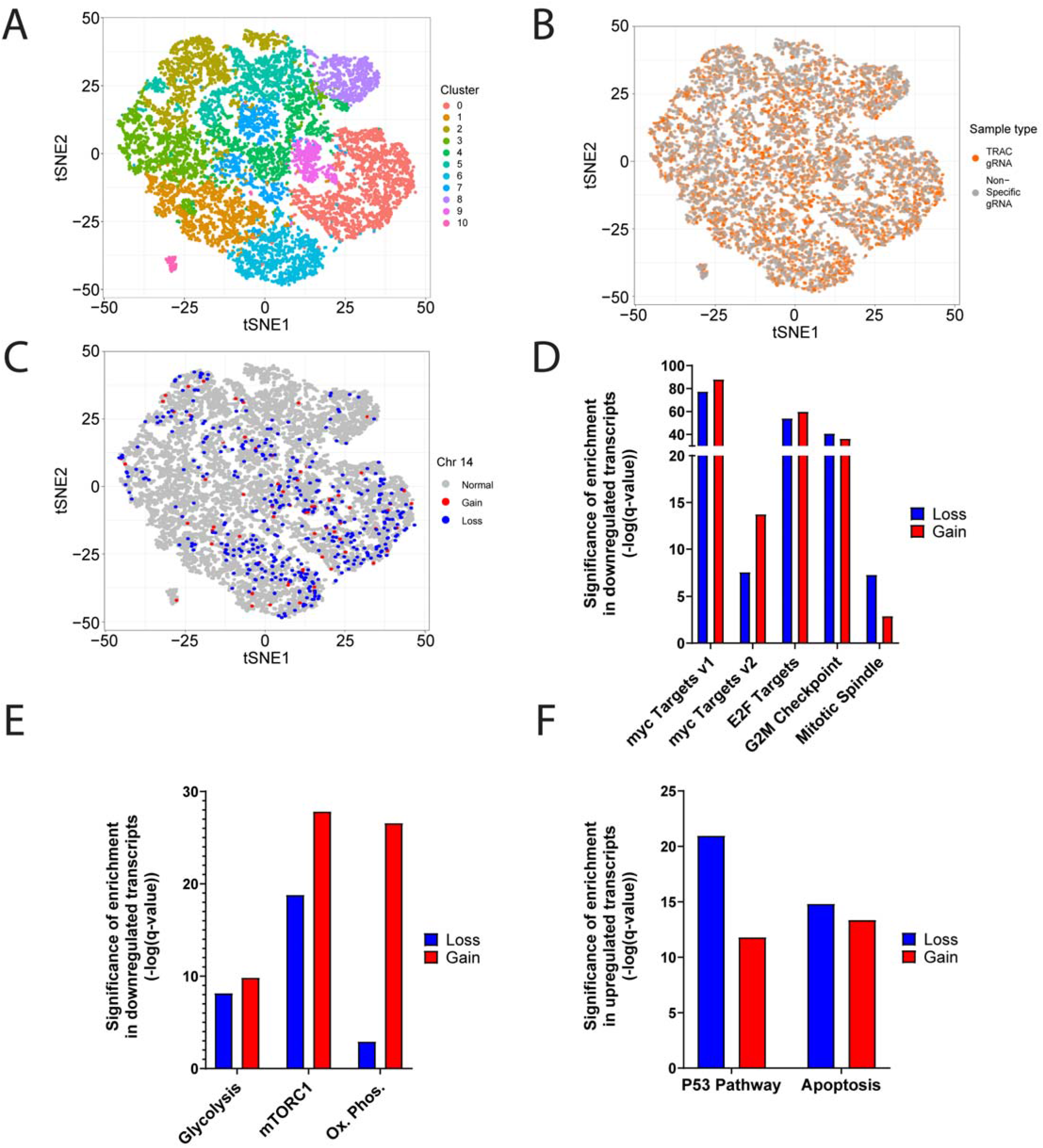
Global gene expression analysis following genome editing. **A**. scRNAseq data from primary human T cells showing 13,755 cells as individual dots. Data is displayed by tSNE with each cluster colored differently. **B**. Cells electroporated with a non-specific (gray) or TCRαtargeting (orange) gRNAs are similarly distributed among the clusters. **C**. Cells with a chromosome 14 loss (blue) or gain (red) are similarly distributed among the clusters. **D-F**. Gene set enrichment analysis reveals that ‘Hallmark’ gene sets associated with cell cycle (D) and metabolism (E) are significantly downregulated; and the ‘Hallmark’ gene sets associated with p53 and apoptosis (F) are significantly upregulated, both in cells with chromosome 14 loss (blue) and in cells with chromosome 14 gain (red). Full enrichment matrix can be found in Extended Data Fig. 5 and Supplementary Table 2.

Targeting different loci concomitantly with several gRNAs may aggravate the risk for chromosomal aberrations. Indeed, Stadtmauer & Fraietta et al. identified multiple translocations between cleaved chromosomes when co-delivering 3 gRNAs targeting both TCR loci as well as the PDCD1 gene^6^. We next used the same gRNA combination for CRISPR-Cas9 RNP electroporation of primary human T cells (Fig. 4A). We verified cleavage efficiency (Extended Data Fig. 6) and performed scRNAseq in order to analyze gene expression patterns and infer copy number changes along the targeted chromosomes. The transcriptional downregulation of TCRα, TCRβ and PDCD1 was confirmed (Fig. 4B). Importantly, in this experiment, we found expression patterns indicative of a chromosome 14 loss and gain in 9.0% and 1.4% of the cells, respectively (Fig. 4C-D, Extended Data Fig. 7). The higher chromosome 14 aberration rate, compared to the previous experiment, may be explained by the 3-fold higher Cas9 concentrations, used in order to preserve the Cas9-per-gRNA ratio in the combined targeting setting. Strikingly, we further found as many as 9.9% of treated cells to be characterized with chromosome 7 truncations, entailing the TCRβ locus (Fig. 4C-D). This result suggests that the repair of nuclease-induced breaks in primary human T cells may be inefficient. Importantly, the extent of the detected aberrations is in good correspondence with the chromosomal position of the cleavage sites. Cleaving the TCRα locus, near the chromosome 14 centromere, leads to loss of the entire arm and to functional whole-chromosome aneuploidy; cleaving the TCRβ locus, in the middle of the chromosome 7 q-arm, leads to the expected truncations and cleaving the PDCD1 gene, which resides near the chromosome 2 q-arm telomere, expectedly has little effect on copy number and gene expression (Extended Data Fig. 8). Finally, a gene set enrichment analysis confirmed the transcriptional differences observed in our first experiment in cells that have lost or gained chromosome 14, and identified very similar pathway enrichments in the cells with a chromosome 7 truncation (Fig. 4E-G, Extended Data Fig. 9 and Supplementary Table 3).

**Figure 4:**
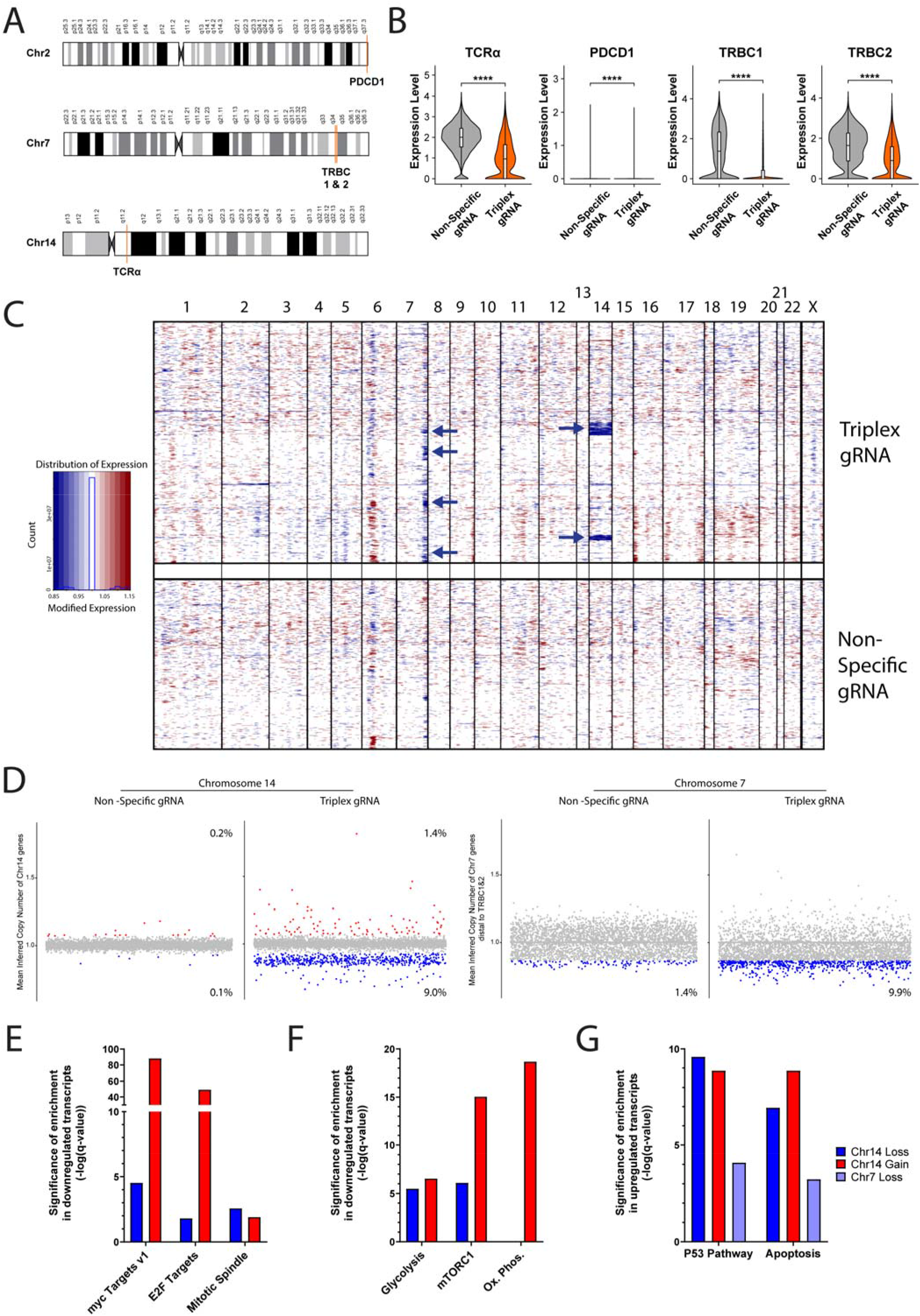
Targeting different loci concomitantly with several gRNAs aggravates chromosomal aberrations. **A**. Schematic depiction of the human chromosomes 2, 7 and 14. Target loci are shown in orange. A single gRNA targets both TRBC1 and TRBC2. **B**. RNA expression of TCRα, PDCD1, TRBC1 and TRBC2 in cells treated with either the non-specific gRNA or the combination of TCRα, TRBCs and PDCD1-targeting gRNAs. ****, p<0.0001, two-sided unpaired Wilcoxon test. **C**. A heat map depicting gene copy numbers inferred from scRNAseq analysis following treatment with either the non-specific or the combination of TCRα, TRBCs and PDCD1-targeting gRNAs. The chromosomes are ordered in columns and the color coding indicates an increase (red) or decrease (blue) in copy number. Blue arrows indicate loss of chromosome 14 or chromosome 7 segments. **D**. Each dot represents the mean inferred copy number of genes coded on the relevant chromosome in each cell treated with a non-specific gRNA (left) the combination of TCRα, TRBCs and PDCD1-targeting gRNAs (right). For chromosome 7, the mean inferred copy number of genes coded distal to the TRBC1&2 is analyzed. Cells are marked with red and blue dots, corresponding to a gain or a loss, respectively, if their mean inferred gene copy number is >2 standard deviations away from the population’s mean. n=8619 and 6326 for cells treated with the non-specific gRNA (left) or the combination of TCRα, TRBCs and PDCD1-targeting gRNAs (right), respectively. **E-G**. Gene set enrichment analysis reveals that ‘Hallmark’ gene sets associated with cell cycle (E) and metabolism (F) are significantly downregulated; and the ‘Hallmark’ gene sets associated with p53 and apoptosis (G) are significantly upregulated, both in cells with chromosome 14 loss (blue) and in cells with chromosome 14 gain (red). Full enrichment matrix can be found in Extended Data Fig. 9 and Supplementary Table 3.

Aneuploidy is associated with most types of cancers^29^. Of note, chromosome 14 loss is frequently found in early-onset colon cancer^30^, and is among the most commonly affected chromosomal regions in ovarian cancer^31^. Chromosome 14 monosomy, trisomy and tetrasomy are also frequently found in meningioma^32^. Similarly, high grade glioma is often characterized by chromosome 7 trisomy^33^. Due to the role of aneuploidy in tumorigenesis, aneuploid cells are considered a tumorigenic risk and are strictly avoided in cell transplantations^34^. Our observations raise the possibility that Cas9-induced aneuploidy and chromosomal truncations, observed in our study, will too be associated with increased risk of tumorigenesis. Importantly, the methods we described were optimized for the detection of aneuploidy and large truncations, therefore underestimating the overall occurrence of genetic alterations. While detected in some cases (Extended Data Fig. 3), translocations and focal deletions may better be assessed using complementary technologies^6^.

Aberration rates can potentially be reduced by using base editors instead of nucleases^35^. In the liver, gene insertion by homologous recombination can be achieved without nucleases^36–38^. However, in T cells, nucleases may still be required for site-specific gene insertion of CAR and engineered TCR genes into TCR loci in order to enhance Tcell potency and delay exhaustion^8–10^. Further studies are required in order to determine whether sorting for expression of the targeted receptor cassette will help reduce the rate of cells with nuclease-induced aberrations. Alternatively, endogenous genes that are differentially expressed in aberrant T cells may be used for cell sorting prior to adoptive transfer. As a first step towards this aim, we compiled a list of genes whose expression significantly differed between the cells that have lost chromosome 14 and those that have not (Supplementary Table 1), and manually curated the list to identify cell surface proteins. Interestingly, CD70, which is expressed at low levels on naïve T cells^39^, is upregulated in cells with a chromosome 14 loss (Extended Data Fig. 10). Similarly, L-Selectin (SELL/CD62L) is highly expressed in naïve T cells^40^ and is downregulated in chromosome 7 truncated cells (Extended Data Fig. 10). Finally, our gene set enrichment analysis suggests that aneuploidy provides fitness disadvantage to the T cells *in vitro* (Fig. 3D-F, Supplementary Table 2-3), in line with multiple studies that showed the negative effect of aneuploidy induction on cell survival and proliferation ^29,41,42^. Additional propagation of the cells in culture prior to transplantation may reduce the prevalence of aneuploid cells, and extended culturing should thus be considered, while also weighing the effects on cell activation, differentiation and exhaustion.

In conclusion, inferring copy number variations from scRNAseq data is an unbiased high-throughput method to identify nuclease-induced chromosomal aberrations. Our discovery of frequent aneuploidy and chromosomal truncations in human T cells targeted using CRISPR-Cas9 and clinical gRNA sequences, highlights potential oncogenic risks and underscores the need for mitigation strategies in order to allow the safe application of nucleases in adoptive T cell transfer and beyond.

## METHODS

### T cell editing

Whole blood was obtained with donor consent from the Israeli Blood Bank (Magen David Adom, Sheiba Medical Center) in accordance with Tel Aviv University Review Board. PBMCs were extracted using Lymphocyte Separation Medium (mpbio) and cryopreserved until subsequent use. Following thawing, cells were activated for 24-48hrs with 1ug/ml anti-human CD28 (biogems) and anti-human CD3 (biogems). Cells were cultured in MEM-Alpha (Biological Industries) supplemented with 10% Heat Inactivated FCS (Sigma), 50IU rhIL-2 (Peprotech) and P/S. For RNP electroporation, 18pmol of Alt-R spCas9 Nuclease V3 (IDT) and 66pmol of Alt-R CRISPR-Cas9 gRNA (IDT) per 1E6 cells per target were complexed in Buffer T (IDT). Experiments involving a triple targets (TRAC, TRBC, PD-1), quantities of Cas9:gRNA complexes were tripled for the non-specific control. gRNA sequences can be found in Supplementary Table 4. Cells were harvested, washed and electroporated at 1600v, 10ms, 3pulses in 10ul Buffer T and subsequently grown in culture media devoid of P/S.

### Flow Cytometry

Cells were harvested, washed, and resuspended in Cell Staining Buffer (Biolegend) containing 1/100 diluted anti-human CD3 or anti-human TCRα/β (Biolegend), both targeting the TCR complex. Staining was performed for 15mins at room temperature in the dark. Finally, cells were washed and data acquisition was performed on an Attune NxT Flow Cytometer (life Technologies).

### Single-cell RNA sequencing

Edited T cells were harvested and live cells were purified using Lymphocyte Separation Medium (mpbio), washed and resuspended in PBS supplemented with 0.5% BSA to achieve optimal concentration of approximately 1000 cells per microliter. Cells were counted and viability assessed manually in Trypan Blue 0.4% (Biological Industries). 17,000 cells were loaded on Next GEM Chip (10x Genomics). Libraries were prepared at the Single-Cell Genomics Core, Faculty of Medicine, Tel Aviv University, using the 10x Genomics Chromium Controller in conjunction with the single-cell 3⍰ v3.1 kit, protocol revision D. The cDNA synthesis, barcoding, and library preparation were then carried out according to the manufacturer’s instructions. Briefly, cDNA amplification was performed for 11 cycles. Sample index PCR was performed for 13 cycles using Chromium i7 Sample Indices. Resulting libraries were quantified and analyzed by Qubit and Tapestation. Libraries were sequenced on the NextSeq 500 platform (Illumina), following the manufacturer’s protocol, using a NextSeq 500/550 High Output Kit v2.5 (75 Cycles) kit (Illumina). Sequencing was performed at the Genomics Research Unit, at the Life Sciences Inter-Departmental Research Facility Unit, Tel-Aviv University.

### scRNA-seq gene expression pre-processing

Raw BCL files for the DNA sequencing data were processed using Cellranger DNA (version 5.0.1). Data were aligned to the 10X Genomics GRCh38 genome. Results were visualized in the Loupe scDNA Browser (version 5.0.0). Raw gene expression data were extracted from the Seurat object as recommended in the “Using 10x data” section (inferCNV of the Trinity CTAT Project, https://github.com/broadinstitute/inferCNV).

### InferCNV

InferCNV was used to infer copy number changes from the gene expression profiles^26,27^ The non-targeted T cell population was used as the reference, and the CRISPR/Cas9-targeted population was tested, with the following parameters: “denoise”, default hidden markov model (HMM) settings, and a value of 0.1 for “cutoff”.

### Identification of aberrant cells

For each cell, the mean of inferCNV scores was calculated across genes and plotted. The PDCD1 gene resides near the chromosome 2 q-arm telomere and only 15 genes, expressed in T cells, reside between the TCRα gene and the chromosome 14 centromere. Therefore, for chromosome 14 and chromosome 2, all of the expressed genes on the respective chromosomes were used for the analysis. For chromosome 7, the 47 expressed genes that reside distal to TCRβ were used. Cells with a mean lower or higher >2 standard deviations from the mean of the population were determined as cells with a loss or a gain, respectively.

### Simulation analysis for downregulated genes

In order to determine whether cells categorized as harboring a chr14 loss of a chr7 distal loss, also had a significant increase in the fraction of zero expression calls (that is, whether these regions are enriched with genes not detected at all by scRNAseq), the ratio between the number of genes from each chromosome with expression = 0 and expression > 0 for each cell population (loss vs. non-loss) was calculated. The fold change of the proportion of zeros calls between the normal and aberrant cells was determined, and 10,000 simulations were then performed, selecting an equivalent number of random genes from other chromosomes. An empirical p-value was determined by comparing the empirical values to the simulated values.

### Differential gene expression analysis

The “FindMarkers” package from the Seurat library^43^ was used to detect the differentially expressed (DE) genes between two groups of cells. The function receives two identities of clusters in the data set and a value for the minimum percentage that is required for a feature to be detected in either of the two groups of cells. The minimum percentage value that we used is 0.25.

The first comparison was made to detect DE genes between the cells that have undergone loss in the TRAC gRNA and the cells that did not show loss or gain in the same treatment group. The second was to detect DE genes between the cell that have undergone gain in the TRAC gRNA and the cells that did not show gain or loss in the same treatment group.

### Gene Set Enrichment Analysis

The lists of differentially expressed genes between each two conditions were determined using Seurat as described above. These lists were subjected to gene set enrichment analysis (GSEA) using the GSEA-MSigDB portal (https://www.gsea-msigdb.org/gsea/msigdb/). The analysis was run using the following curated gene sets: “Hallmark”, “KEGG”, “GO biological process” and “positional” gene sets from MSigDB^44–46^.

### ddPCR

For digital droplet PCR, whole genomic DNA was extracted from cells using Gentra PureGene Tissue Kit (Qiagen). In order to remove sheared genomic fragments, resulting eluates were further purified using AmpureXP beads (Beckman Coulter) at a 0.5:1 ratio. DNA fragmentation by digestion was performed in reaction, using 66ng of purified genomic DNA and 10U HindIII-HF (NEB) in ddPCR Supermix for Probes (BioRad). Thermo-cycling reaction was performed as per manufacturer recommendation. Sequences for the primers and probes can be found in Supplementary Table 4. Reactions were performed using a QX200 Droplet Digital PCR System (Bio-Rad). To analyze for dislinkage, we used the following equation as per the resulting Quantasoft (BioRad) Linkage (*Linkage*) and Concentration (C_*HEX*_, C_*FAM*_) values:

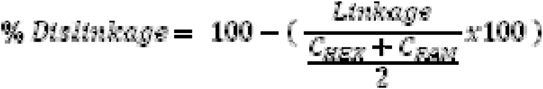

For dislinkage followup, multiple electroporations of treated cells were pooled and then divided in separate wells for collection at each time point. Cells were seeded at 1E6 cells/ml in MEM-Alpha (Biological Industries) supplemented with 10% Heat Inactivated FCS (Sigma) and 50IU rhIL-2 (Peprotech).

### Nucleic acid manipulations

For T7E1 assays, PCR amplification was performed on Gentra PureGene Tissue Kit (Qiagen) extracted genomic DNA. 200-500ng of genomic DNA was amplified using PrimeStar MAX (Takara) for 35 cycles. Primers for these reactions can be found in Supplementary Table 4. Resulting amplicons were denatured and reannealed in a thermocycler before nuclease reaction using T7 Endonuclease 1 (New England Biolabs) at 37C for 20min. Resulting fragments were analyzed by agarose gel electrophoresis and quantified using Biovision (Vilber Lourmat) using a rolling ball for background subtraction. Efficiency was calculated using the following equation:

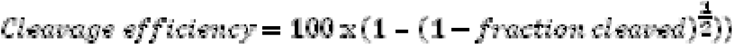

For TIDE analysis, PCR amplicons were subjected to purification by AmpureXP beads (Beckman Coulter) at a 1:1 ratio. Sanger sequencing was performed at the DNA Sequencing Unit, Tel Aviv University. Samples were compared using TIDE (https://tide.nki.nl/)^47^.

### Fluorescence in-situ hybridization

Fluorescence in situ hybridization (FISH) analysis was performed following the manufacturer’s instructions (Cytocell) on interphase human T cells from peripheral blood spreads’ using the TRACD breakapart probe. Images were captured using GenASIs imaging system.

### Statistics

For FISH, genomic DNA cleavage efficiency and flow cytometry knock out efficiency, statistical analyses were performed on Prism (GraphPad). For t-tests on dislinkage by ddPCR, for each donor, technical replicates were averaged and t-test was performed on the averaged values. Each figure legend denotes the statistic used, central tendency and error bars.

## ACKOWLEDGEMENTS

We thank Natalie Zelikson (TAU), for helpful discussions and reviewing the manuscript. We thank the IDRFU, GRU and SICF units, Tel Aviv University for logistic support and council. This research was funded by the European Research Council (A.B. and U.B.-D.), the Gertner Institute Scholarship, the Yoran Institute Scholarship, the SAIA Foundation (A.D.N.), SCGC Tel Aviv University (A.B. and U.B.-D.), the Israel Cancer Research Fund Gesher Award (U.B.-D.), the Azrieli Faculty Fellowship (U.B.-D), the Alon Fellowship for Outstanding Young Scientists, Israel Council for Higher Education (A.M).

## AUTHOR CONTRIBUTIONS

A.D.N. designed, performed and analyzed the experiments; E.R. and E.G. analyzed the single-cell RNA sequencing data; T.T. and M.L. performed fluorescence in-situ hybridization; M.H-F. helped with sample culturing and processing; R.K. and H.K. performed single-cell RNA sequencing; E.R. supervised fluorescence in-situ hybridization; A.M. and U.B.-D. supervised single-cell RNA sequencing analysis; A.D.N. and A.B. drafted the manuscript, and revised it together with A.M. and U.B.-D; U.B.-D. and A.B. supervised the study; A.B. conceptualized the study.

## CODE AVAILABILITY

The software used for individual and integrated analyses are described and referenced in the individual sections in Methods.

## DATA AVAILABILITY

All datasets are available within the article and its Supplementary Information. 10X scRNAseq data have been deposited to SRA with BioProject accession number PRJNA745549.

Supplementary Information and Extended Data Figures are available for this paper.

## EXTENDED DATA FIGURES

**Extended Data Figure 1:**
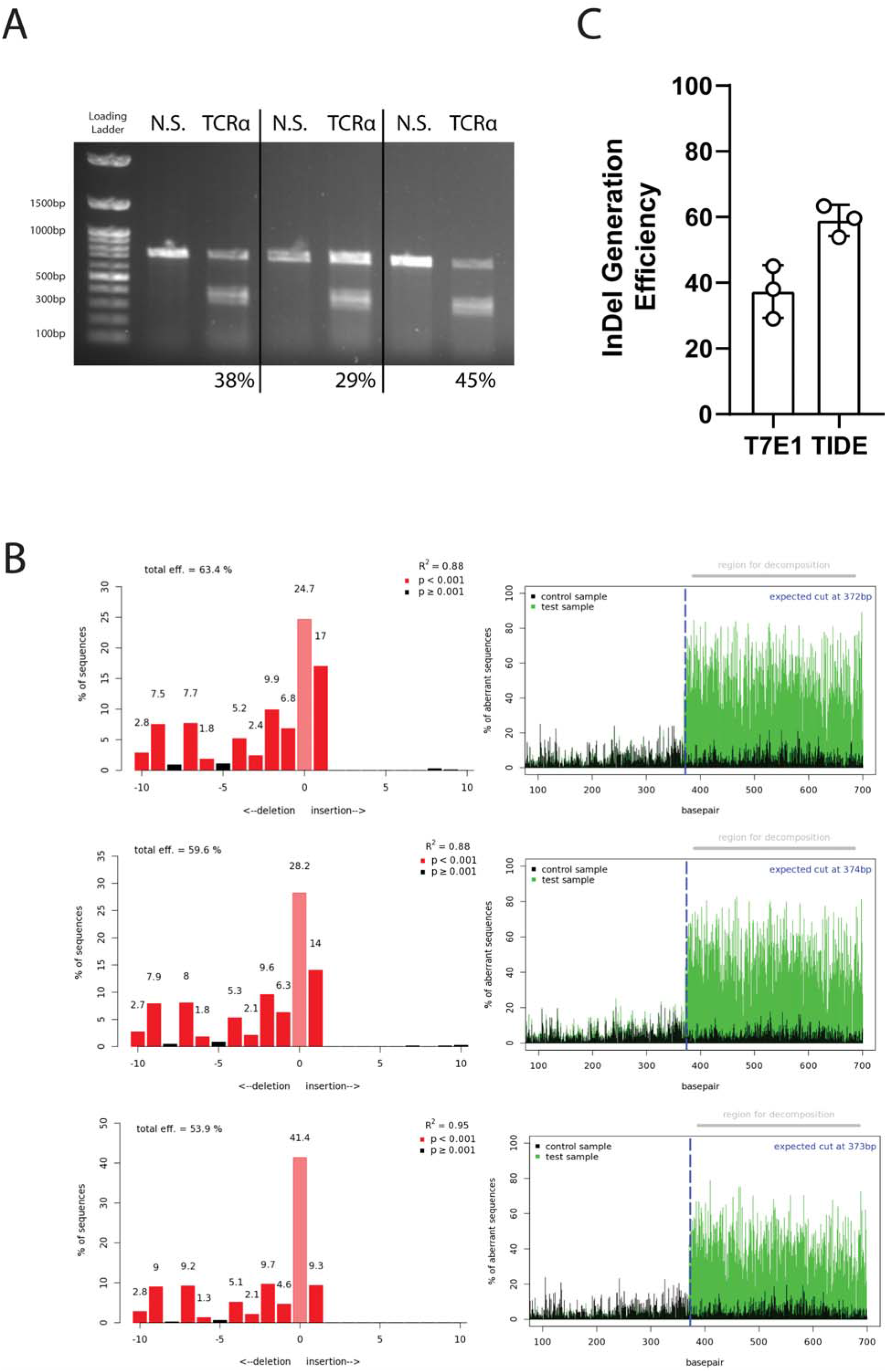
Quantification of InDels produced by CRISPR-Cas9 activity at the TCRα locus indicates efficient cleavage. **A**. T7 Endonuclease 1 (T7E1) assay for three independent experiments. For each experiment, one lane for Non-Specific gRNA (N.S.) and one lane for TCRα-targeting gRNA treated cells are presented. Experiments are separated by a black bar. Percentages, presented below the lanes, refer to cleavage efficiency as inferred by densitometric analysis. Loading Ladder and relative sizes are indicated on the left. Unprocessed scan can be found in Supplementary Data 1. **B**. TIDE Analysis for the same experiments as in A. In the left panel, the height of each bar corresponds to the rate of sequences having the given number of nucleotides added or deleted. The right panel depicts the rate of sequence misalignments at each position of the PCR fragment amplified from the TCRα locus of cells treated with either the TCRα-targeting gRNA (green) or a Non-Specific gRNA (black) **C**. Quantification of A and B.

**Extended Data Figure 2:**
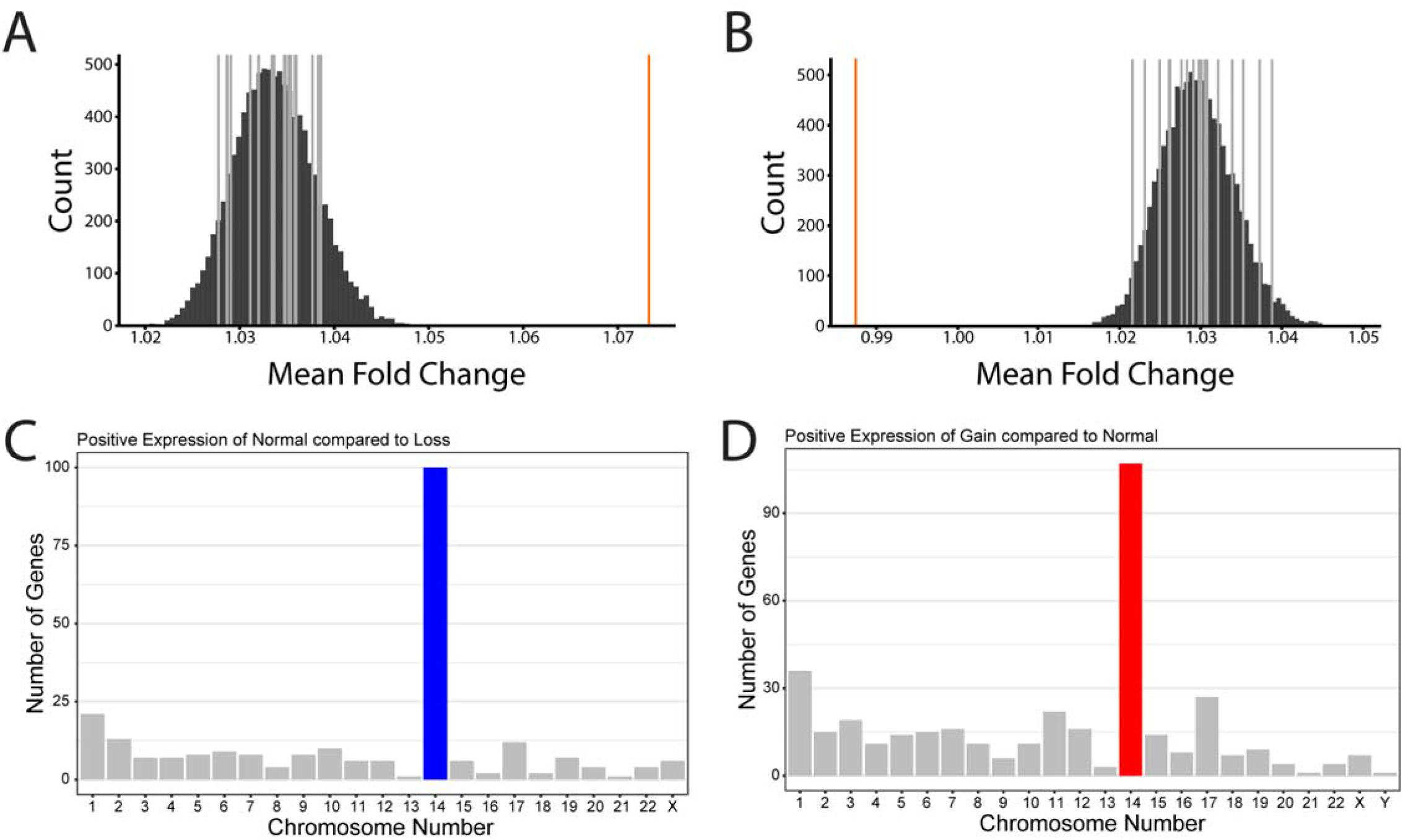
**A-B**. Enrichment in the number of genes with no detected expression among cells identified as having a chromosome 14 loss (A) or gain (B) (Fig. 1F). The x-axis represents the fold-change in the number of genes with no detected expression between cells with or without a chromosome 14 loss, based on the InferCNV analysis (Fig. 1F). The dark gray lines represent the empirical values obtained for each chromosome, except for chromosome 14. The orange line is the empirical value for chromosome 14. The black bars are the results of 10,000 permutations. **C**. Number of differentially expressed genes in each chromosome, as compared between the chromosome 14 loss populations to chromosome 14 normal (see Fig. 1F). **D**. Number of differentially expressed genes in each chromosome, as compared between the chromosome 14 gain populations to chromosome 14 normal (see Fig. 1F).

**Extended Data Figure 3:**
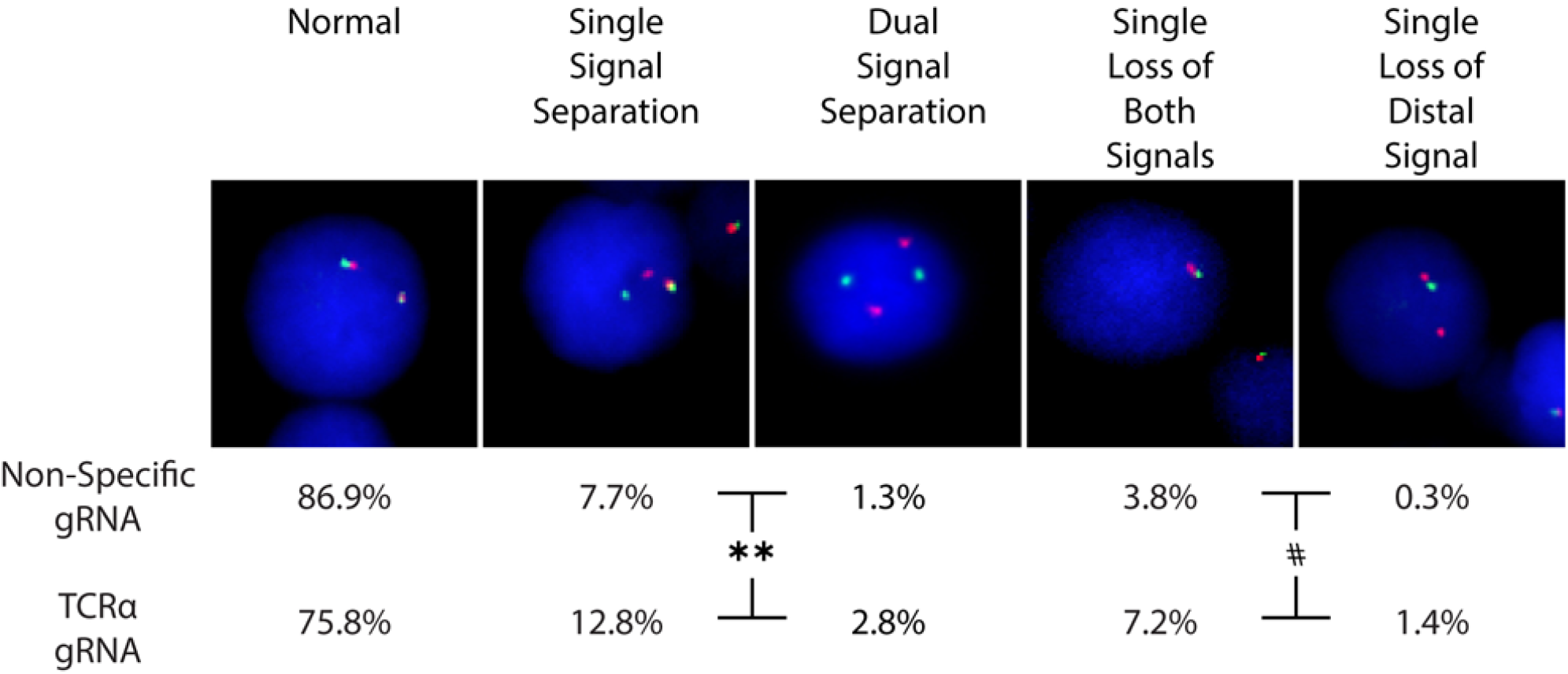
FISH analysis of cells treated with either a non-specific or a TCRα-targeting gRNA. In the pictures are examples of all signal patterns found among analyzed cells. The frequency of each signal pattern, among cells electroporated with the TCRα-targeting gRNA or a non-specific gRNA, is presented below the pictures. n = 313 and n = 360 for non-specific and TCRα targeted cells, respectively. #, p= 0.0115 Fisher’s exact test for loss of either both signals or the distal signal. ** p =0.0048 for either single or dual Signal Separation.

**Extended Data Figure 4:**
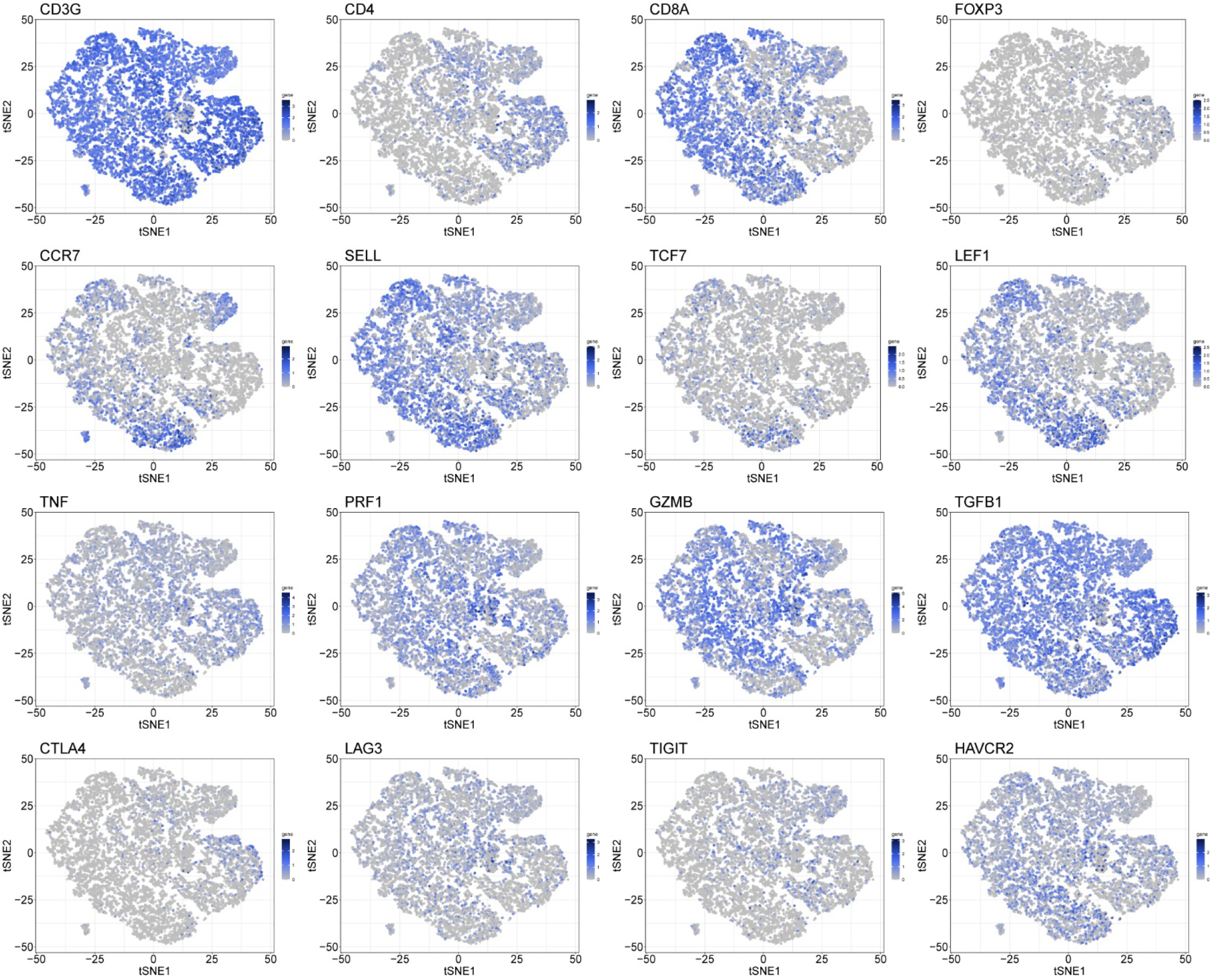
Selected gene expression patterns across the cells. Each t-SNE plots represent all the cells in the experiment (Fig. 3A-C). Each plot represents the expression pattern of a different gene, indicated on the top left, among the clusters. A darker shade of blue corresponds to higher RNA expression.

**Extended Data Figure 5:**
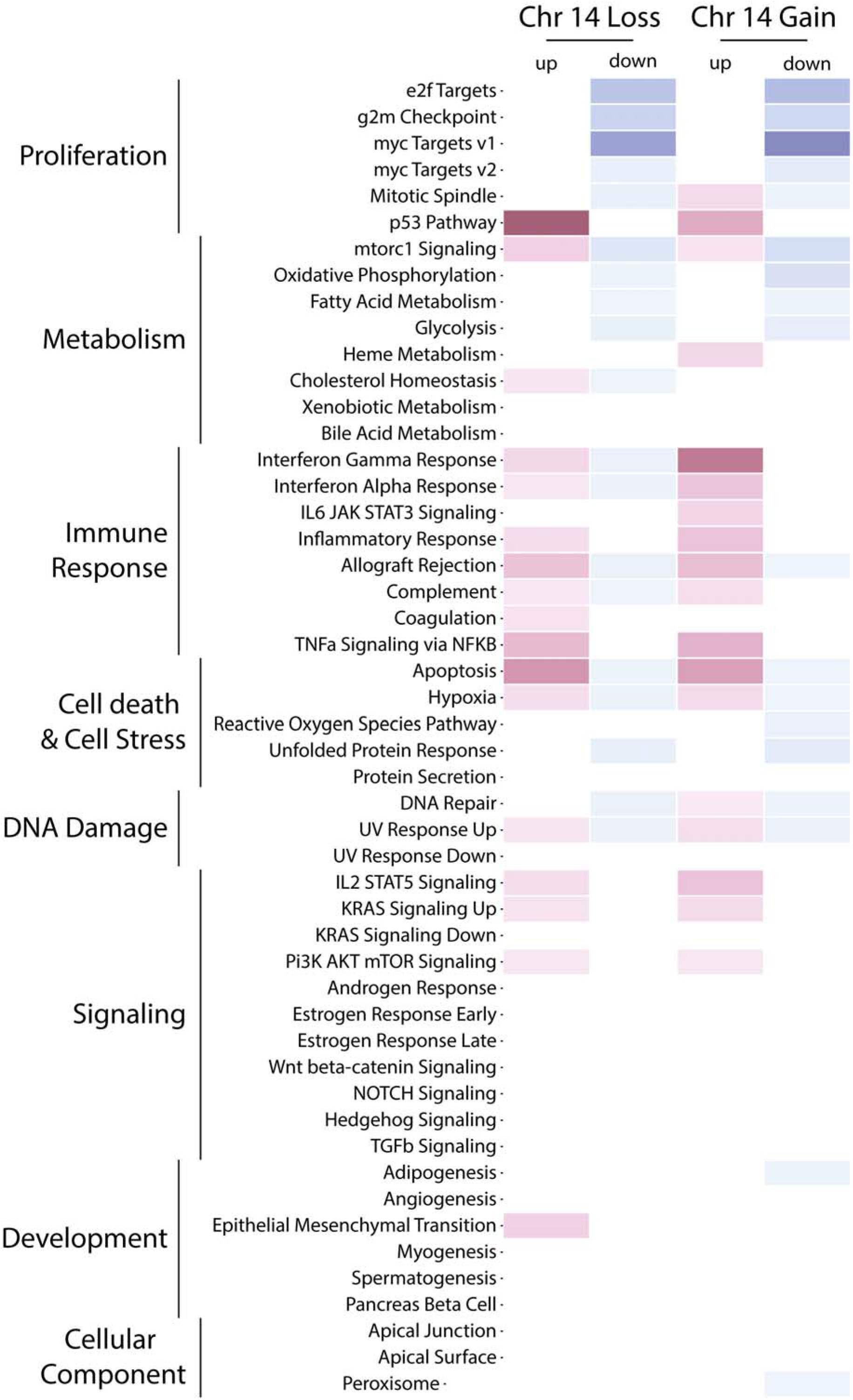
A gene set enrichment analysis between cells that have lost a copy of chromosome 14 (Fig. 1F), to cells without these chromosomal aberrations. The 50 ‘Hallmark’ MSigDB gene sets are shown. Gene sets enriched in up-regulated genes are depicted in red, and those enriched in down-regulated genes are depicted in blue. Values are scaled to -log(FDR) of the enrichments. The full enrichment scores are shown in Supplementary Table 2.

**Extended Data Figure 6:**
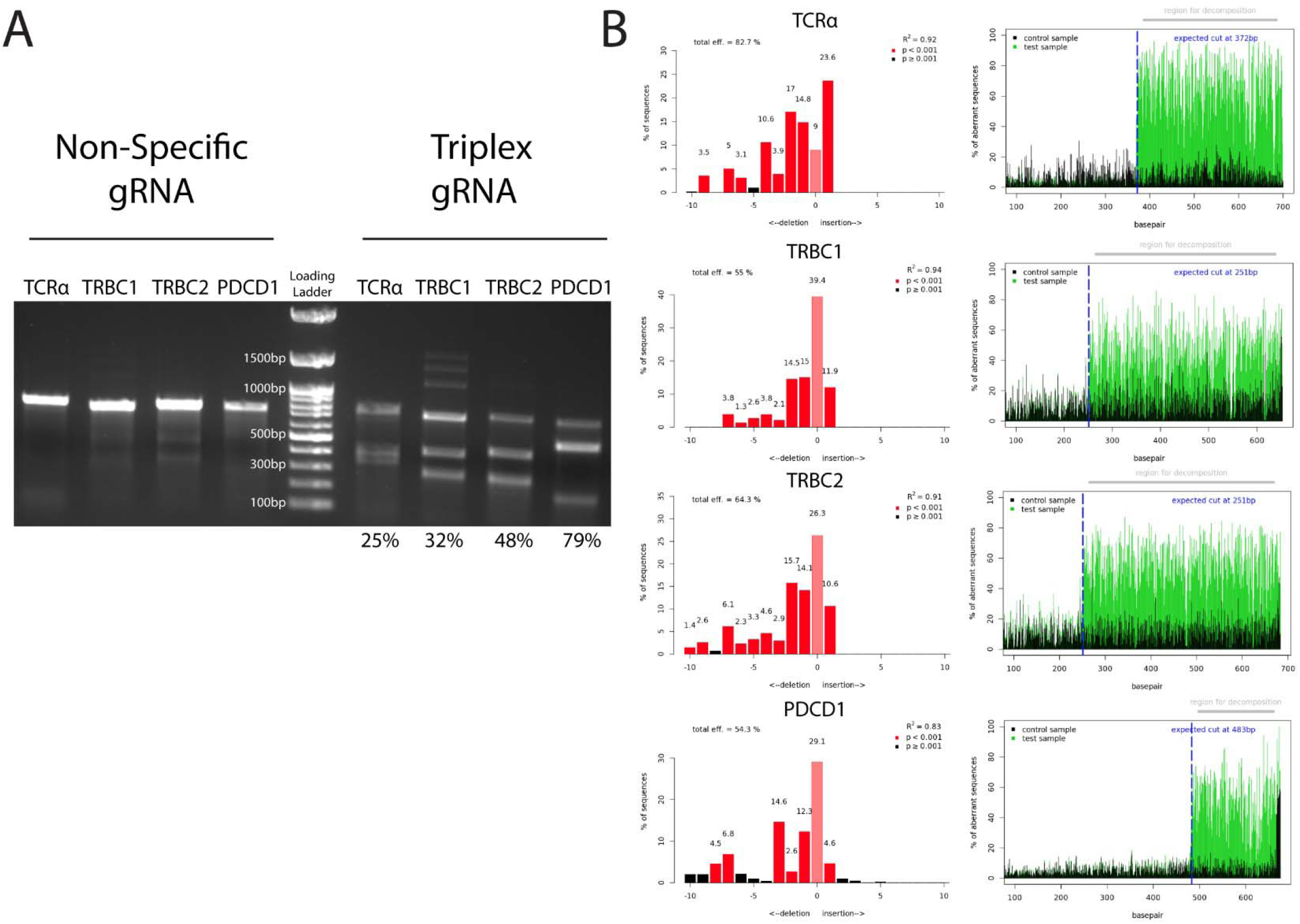
Quantification of InDels produced by CRISPR-Cas9 activity at the TCRα, TCRβ and PDCD1 loci indicates efficient cleavage. **A**. T7E1 assay for the 4 genomic target loci. Non-Specific gRNA treated cells (4 left lanes) are compared to cells treated with the combination of the TCRα, TCRβ and PDCD1-targeting gRNAs (4 right lanes). In each lane, a different locus is analyzed. Calculated efficiency is indicated below. Loading Ladder and relative sizes are indicated in the middle. Unprocessed scan can be found in Supplementary Data 1. **B**. TIDE Analysis for the same experiments as in A. The rows represent (in this order, from top to bottom) the TCRα, TRBC1, TRBC2 and PDCD1 locus. In the left panels, the height of each bar corresponds to the rate of sequences having the given number of nucleotides added or deleted. The right panels depict the rates of sequence misalignments at each position of the PCR fragment amplified from the target locus of cells treated with either Non-Specific gRNA (black) or the combination of the TCRα, TCRβ and PDCD1-targeting gRNAs (green).

**Extended Data Figure 7:**
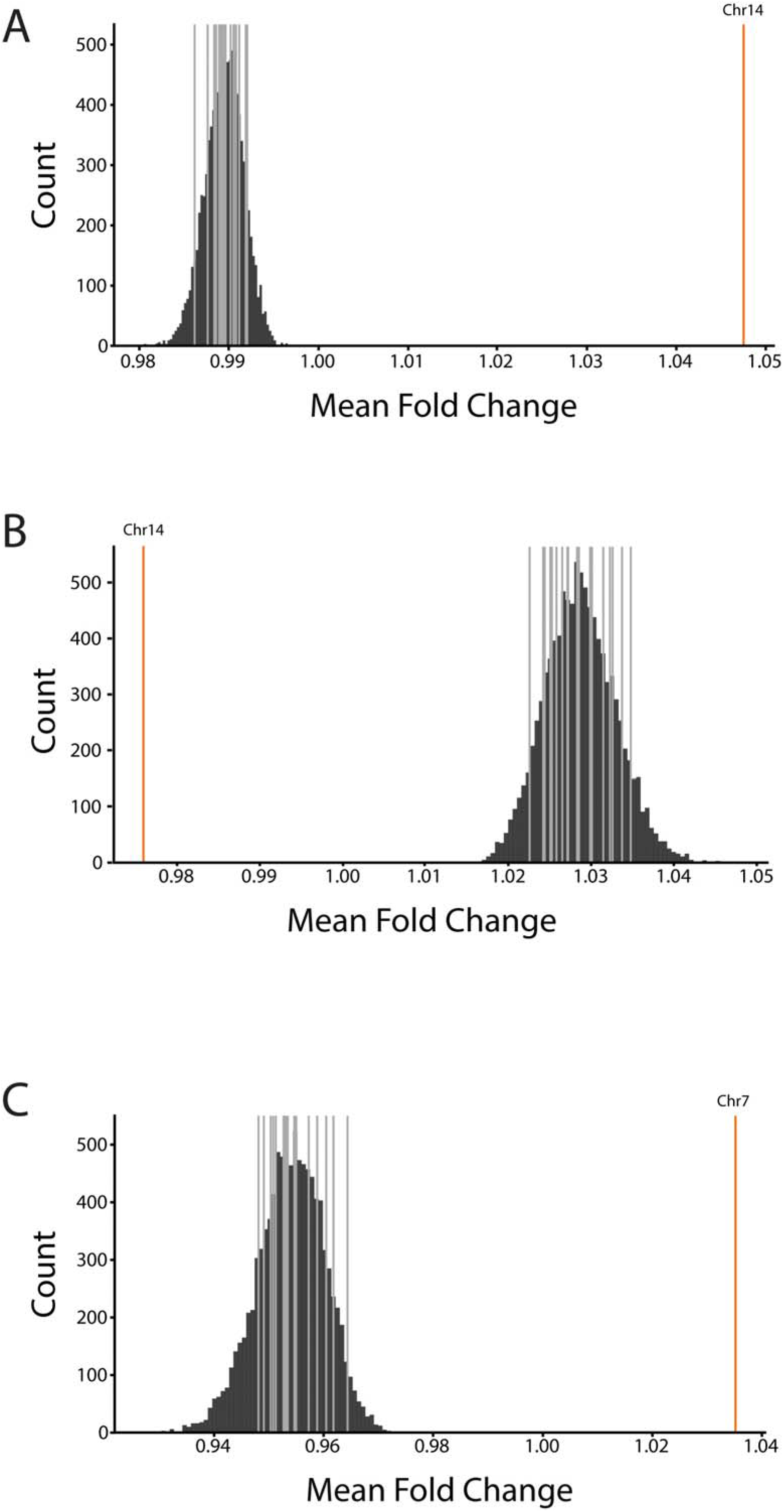
**A-C**. Enrichment in the number of genes with no detected expression among cells identified as having a chromosome 14 loss (A), a chromosome 14 gain (B) or chromosome 7 loss (C) (Fig.4D). The x-axis represents the fold-change in the number of genes with no detected expression between cells with or without a chromosome 14 loss or gain or 7 loss, based on the InferCNV analysis (Fig. 4D). The dark gray lines represent the empirical values obtained for each chromosome, except for chromosome 14 or 7. The orange line is the empirical value for chromosome 14 or chromosome 7. The black bars are the results of 10,000 permutations.

**Extended Data Figure 8:**
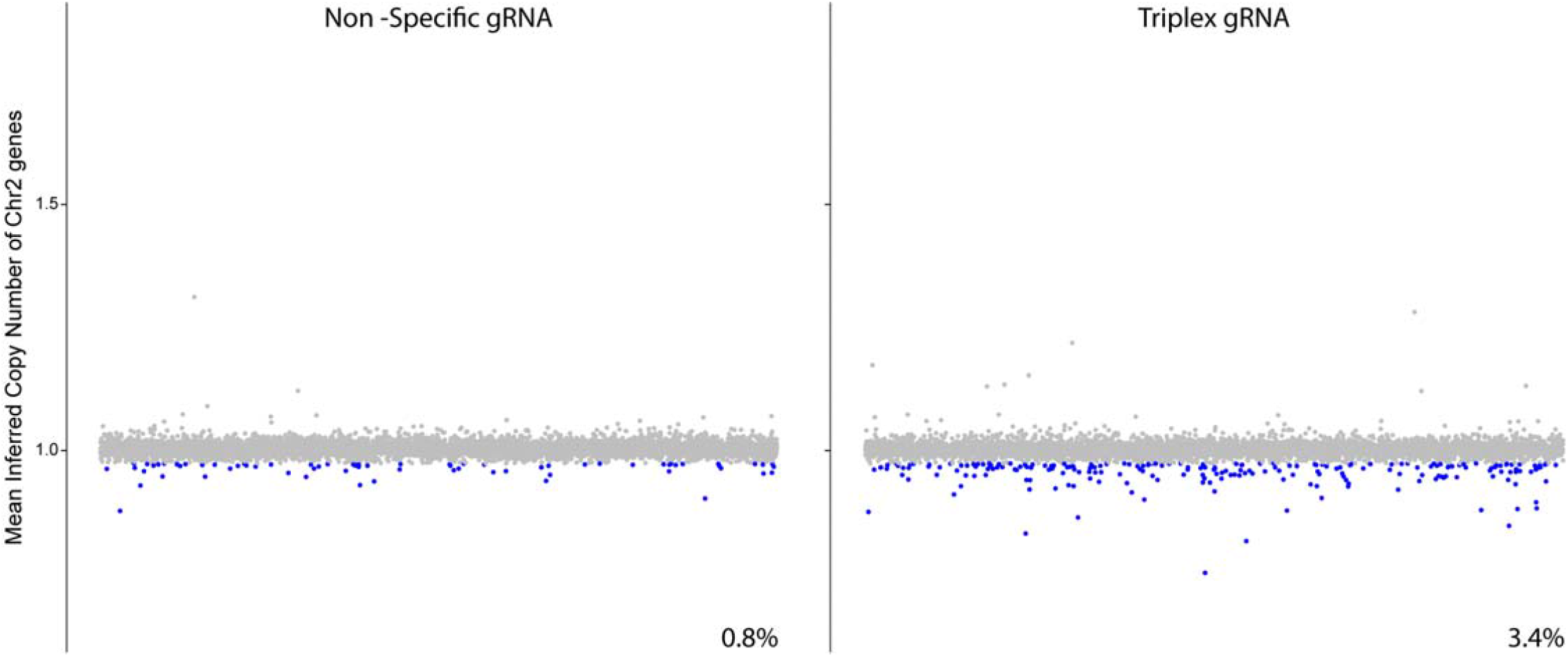
Differential gene expression analysis for chromosome 2. Each dot represents the mean inferred copy number of genes coded on chromosome 2 in each cell treated with a non-specific gRNA (left) or the combination of the TCRα, TCRβ and PDCD1-targeting gRNAs (right). Cells are marked with red and blue dots, corresponding to a chromosome 2 gain or loss respectively, if their mean inferred gene copy number is >2 standard deviations from the population’s mean. n=8619 and 6326 for cells treated with the non-specific gRNA (left) or the combination of the TCRα, TCRβ and PDCD1-targeting gRNAs (right), respectively.

**Extended Data Figure 9:**
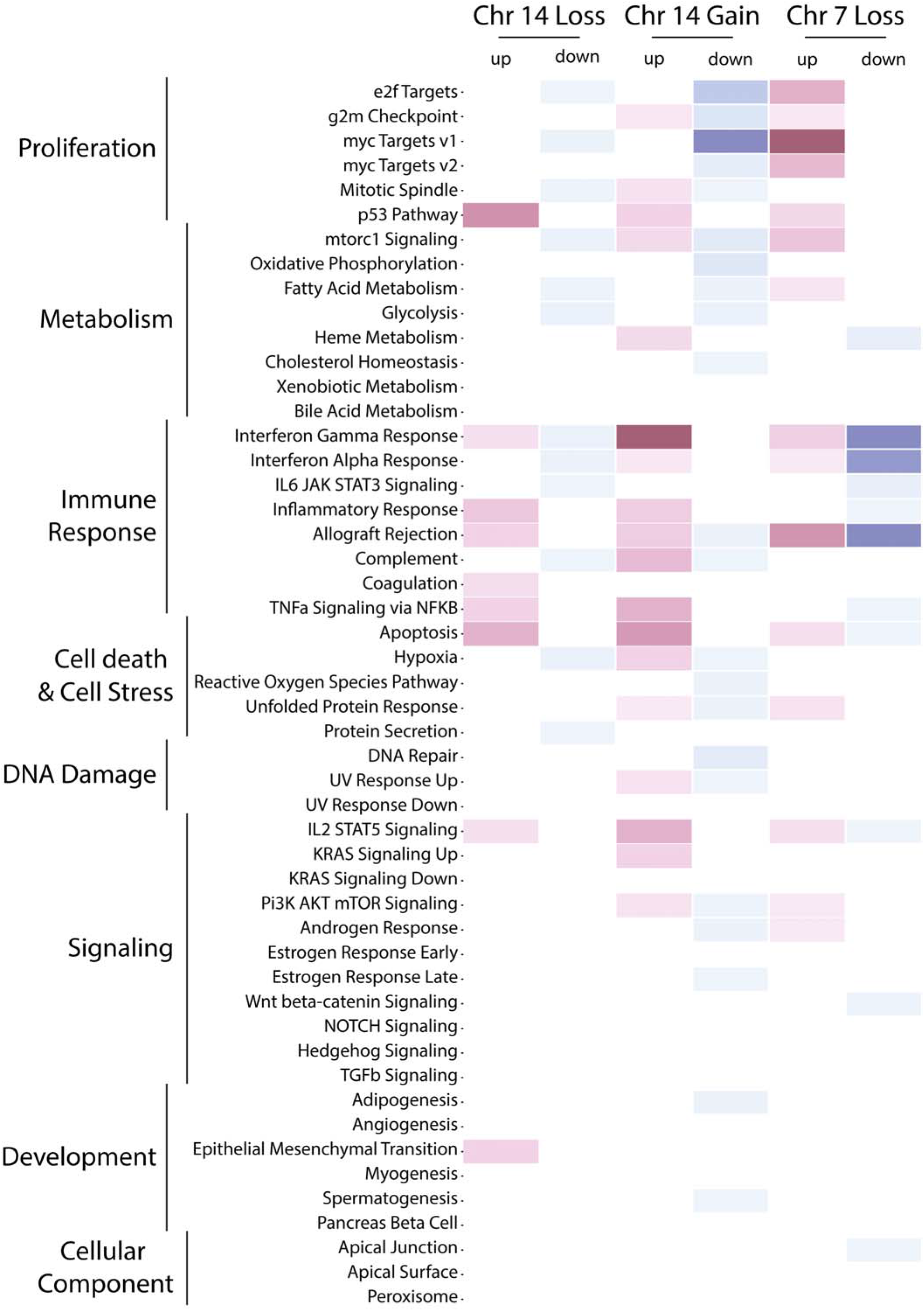
A gene set enrichment analysis between cells that have lost a copy of chromosome 14, or a segment of chromosome 7, to cells without these chromosomal aberrations. The 50 ‘Hallmark’ MSigDB gene sets are shown. Gene sets enriched in up-regulated genes are depicted in red, and those enriched in down-regulated genes are depicted in blue. Values are scaled to -log(FDR) of the enrichments. The full enrichment scores are shown in Supplementary Table 3.

**Extended Data Figure 10:**
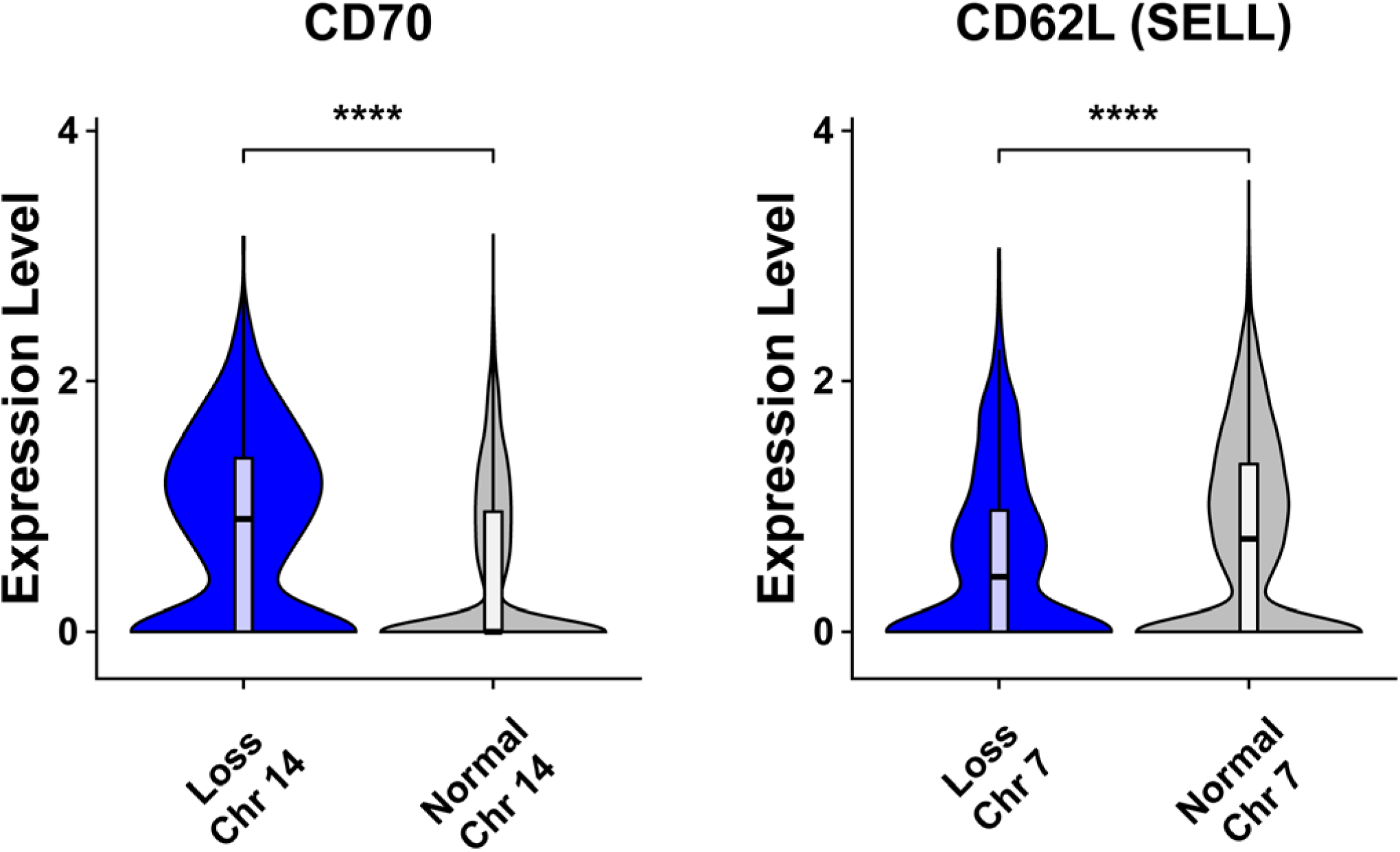
T cell surface markers CD70 and CD62 can potentially be used to sort out aberrant cells. The violin plots show gene expression of CD70 (left) and CD60L (SELL, right), for cells treated with the combination of the TCRα, TCRβ and PDCD1-targeting gRNAs and have normal gene expression (grey) or a gene expression pattern indicating a chromosome 14 loss or a chromosome 7 truncation (blue). **** p<0.0001, unpaired two-tailed Wilcoxon test.

